## Supplementary Figures for "Expansion microsopy reveals *Plasmodium falciparum* blood-stage parasites undergo anaphase with a chromatin bridge in the absence of mini-chromosome maintenance complex binding protein"

SUPPLEMENTARY INFORMATION

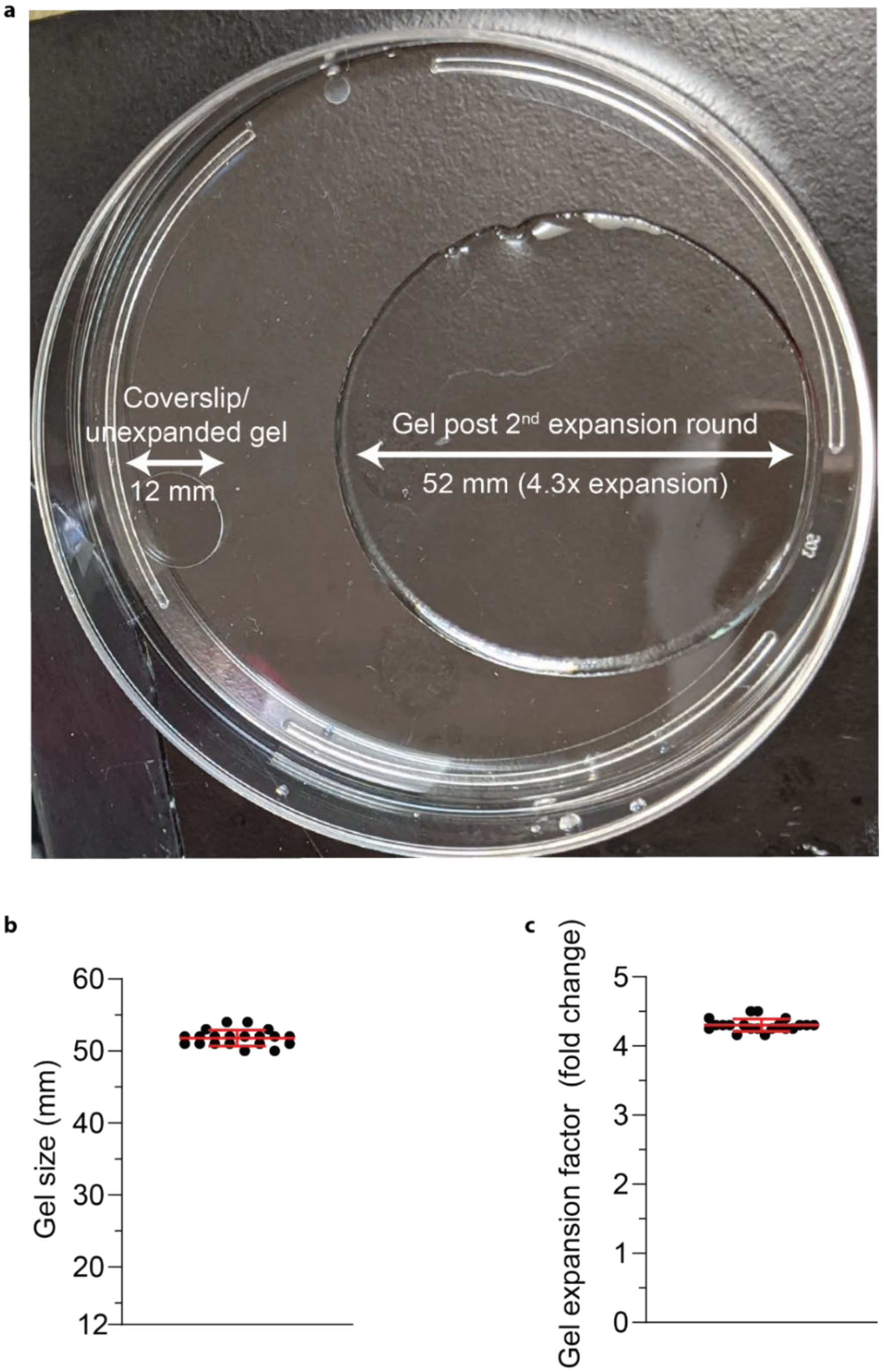


**Supplementary Figure 1. Measurement of U-ExM gels used in this study.**
***(a)*** *Photograph of a 12 mm θ coverslip, on which gelation occurs, next to a fully expanded gel.* ***(b)*** *Final expanded size of all U-ExM gels used in this study, and* ***(c)*** *their expansion factor. n = 20 gels across 4 biological replicates. Error bars = SD.*

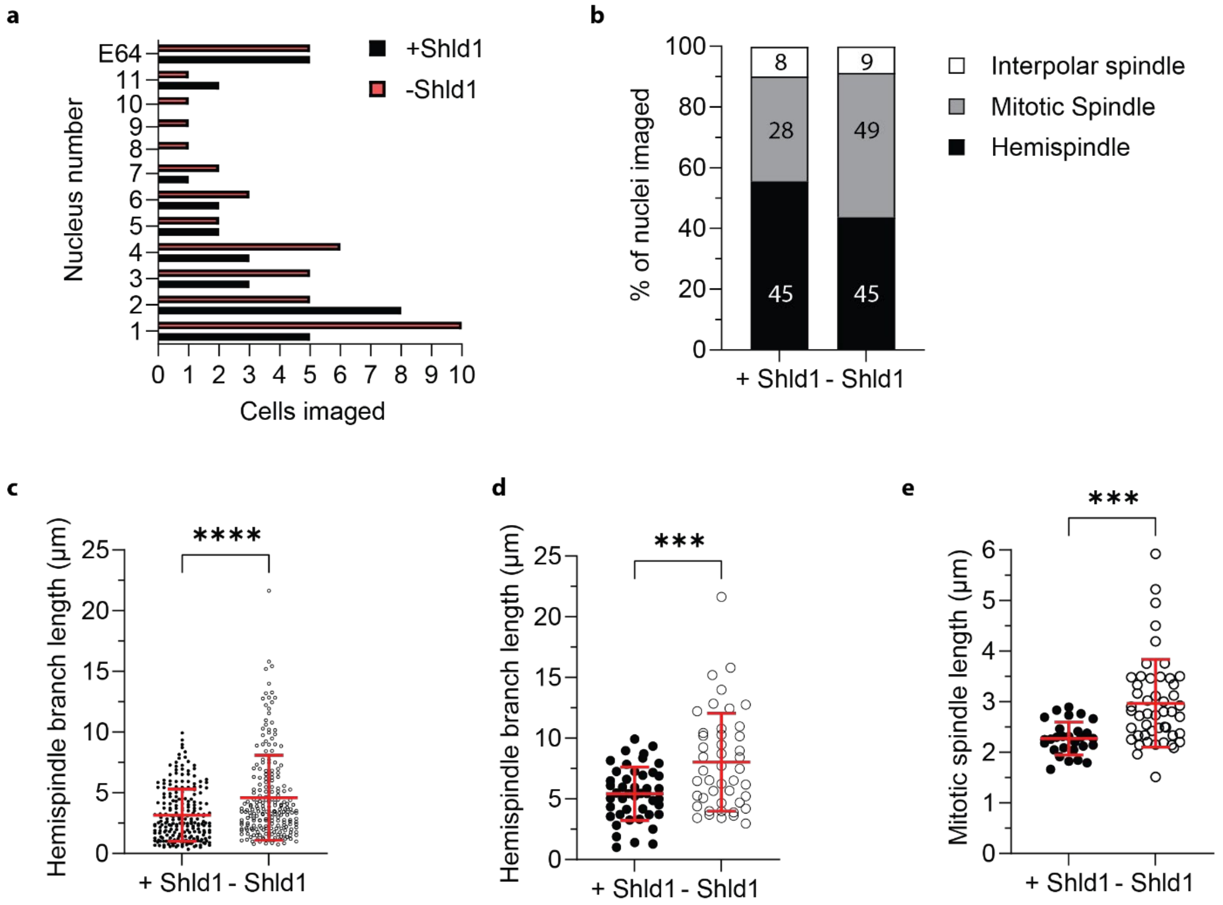


**Supplementary Figure 2. Parasite population summary, spindle type summary and expanded measurements.**
***(a)*** *Histogram of the number of nuclei in each pre-segmentation cell imaged with U-ExM for either* $MCMBP^{HADD}$ *+Shld1 (black) or -Shld1 (red) parasites in this study. E64 = E64-treated, segmented schizonts.* ***(b)*** *Summary of microtubule types observed in imaged nuclei from this study as a % of total nuclei image. Number of each microtubule structure imaged displayed in the corresponding section of the bar graph. Parasites where subpellicular microtubules were observed are not included in this summary. Raw measurements from expanded parasites of* ***(c)*** *individual hemispindle branch length,* ***(d)*** *longest branch in each hemispindle, and* ***(e)*** *mitotic spindles. Estimated actual measurements from these data are presented in Figure 2 (*** = $p < 0.001$, **** = $p < 0.0001$ by unpaired two-tailed t-test, error bars = SD).*

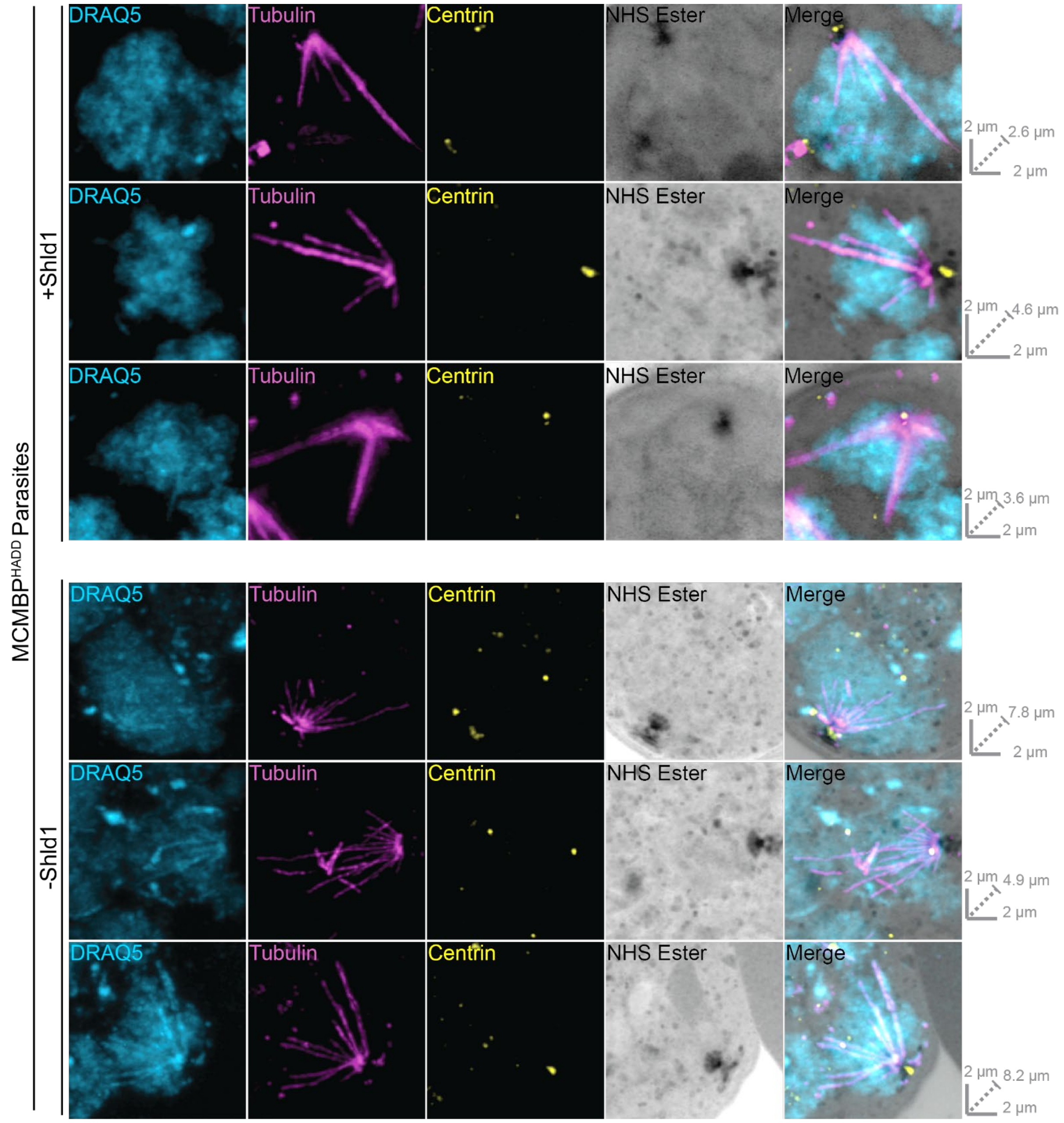


**Supplementary Figure 3. Examples of hemispindles in MCMBP$^{HADD}$ parasites in the presence and absence of Shld1.**

*Representative Airyscan microscopy images of hemispindles from MCMBP$^{HADD}$ parasites cultured either [+]/[-] Shld1 and prepared for U-ExM after staining with a nuclear stain (DRAQ5, in cyan), anti-tubulin (in magenta), anti-centrin (in yellow), and a protein stain (NHS Ester, in grayscale. All images are maximum intensity projections. Scale bars as labelled in each image, solid bars = XY scale, dashed bar = combined depth of slices used for Z-projection.*

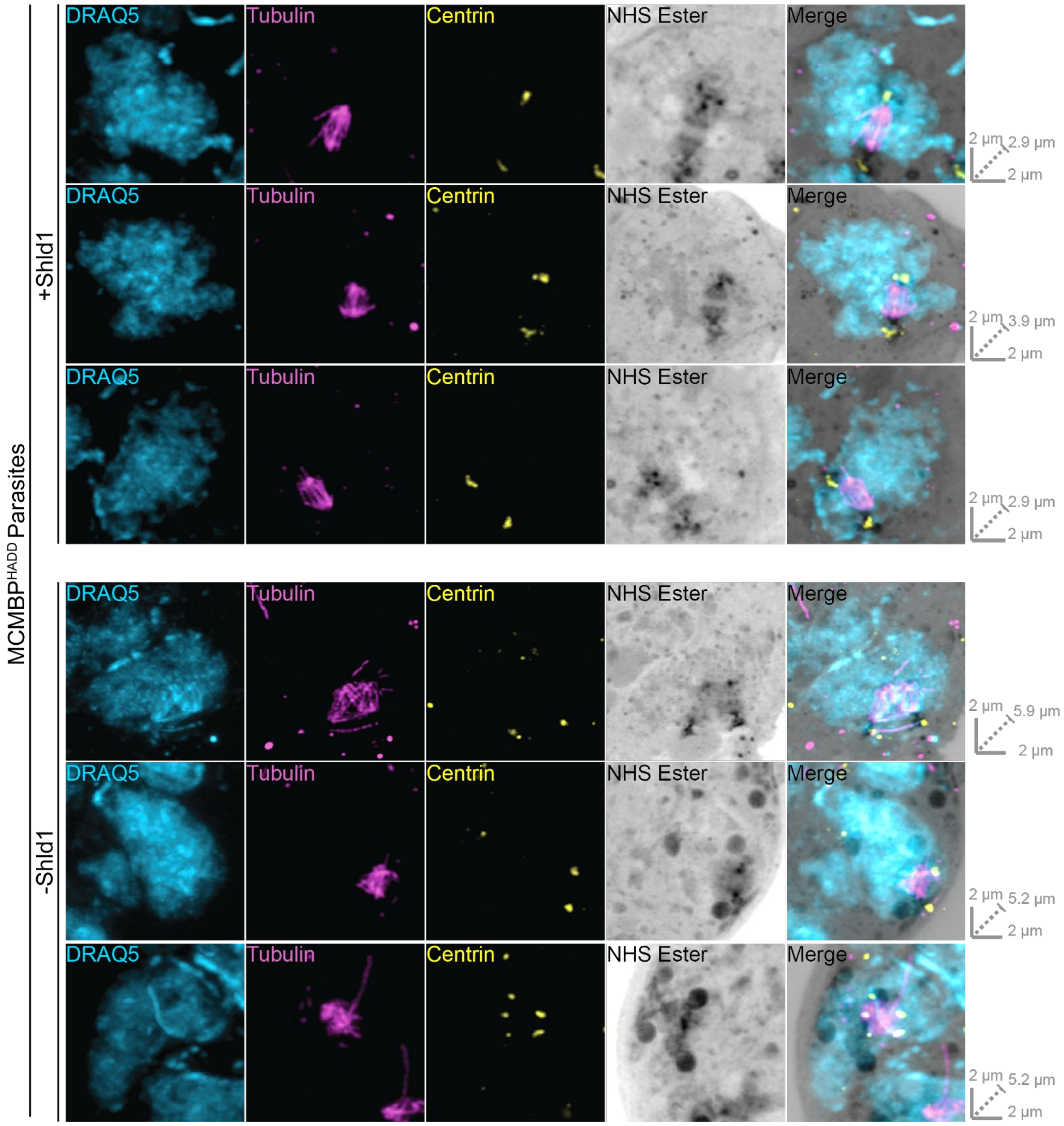


**Supplementary Figure 4. Examples of mitotic spindles in MCMBP$^{HADD}$ parasites in the presence and absence of Shld1.**

*Representative Airyscan microscopy images of mitotic from MCMBP$^{HADD}$ parasites cultured either [+]/[-] Shld1 and prepared for U-ExM after staining with a nuclear stain (DRAQ5, in cyan), anti-tubulin (in magenta), anti-centrin (in yellow), and a protein stain (NHS Ester, in grayscale. All images are maximum intensity projections. Scale bars as labelled in each image, solid bars = XY scale, dashed bar = combined depth of slices used for Z-projection.*

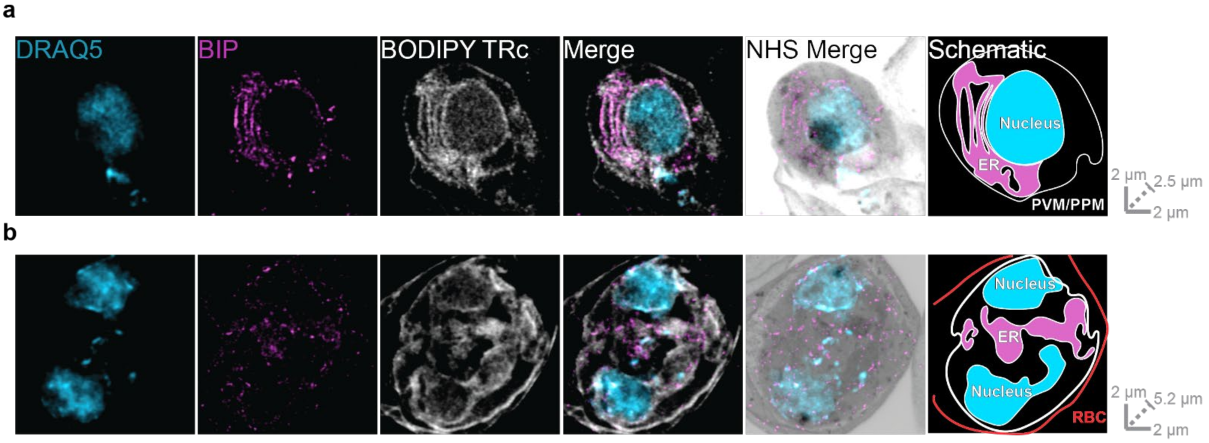


**Supplementary Figure 5. BODIPY TRc stains the PPM, PVM, ER, nuclear envelope and RBC membrane.** *3D7 parasites were then prepared for U-ExM, stained with a nuclear stain (DRAQ5, in cyan), anti-BIP (ER marker, in magenta), a membrane stain (BODIPY TRc, in white), and a protein stain (NHS Ester, in grayscale), and visualized using Airyscan microscopy.* ***(a)*** *A ring stage parasite shows partial colocalization between BODIPY TRc and BIP, as well showing BODIPY TRc staining encircling both the nucleus and the parasite plasma membrane (PPM) and/or parasite vacuole membrane (PVM); which cannot be differentiated in this image.* ***(b)*** *A two nucleus trophozoite shows a similar staining pattern, although the RBC membrane, which is also stained by BODIPY TRc, can also be observed. Images containing BODIPY TRc are average intensity projections, while those with NHS ester are maximum intensity projections. Scale bars as labelled in each image, solid bars = XY scale, dashed bar = combined depth of slices used for Z-projection.*

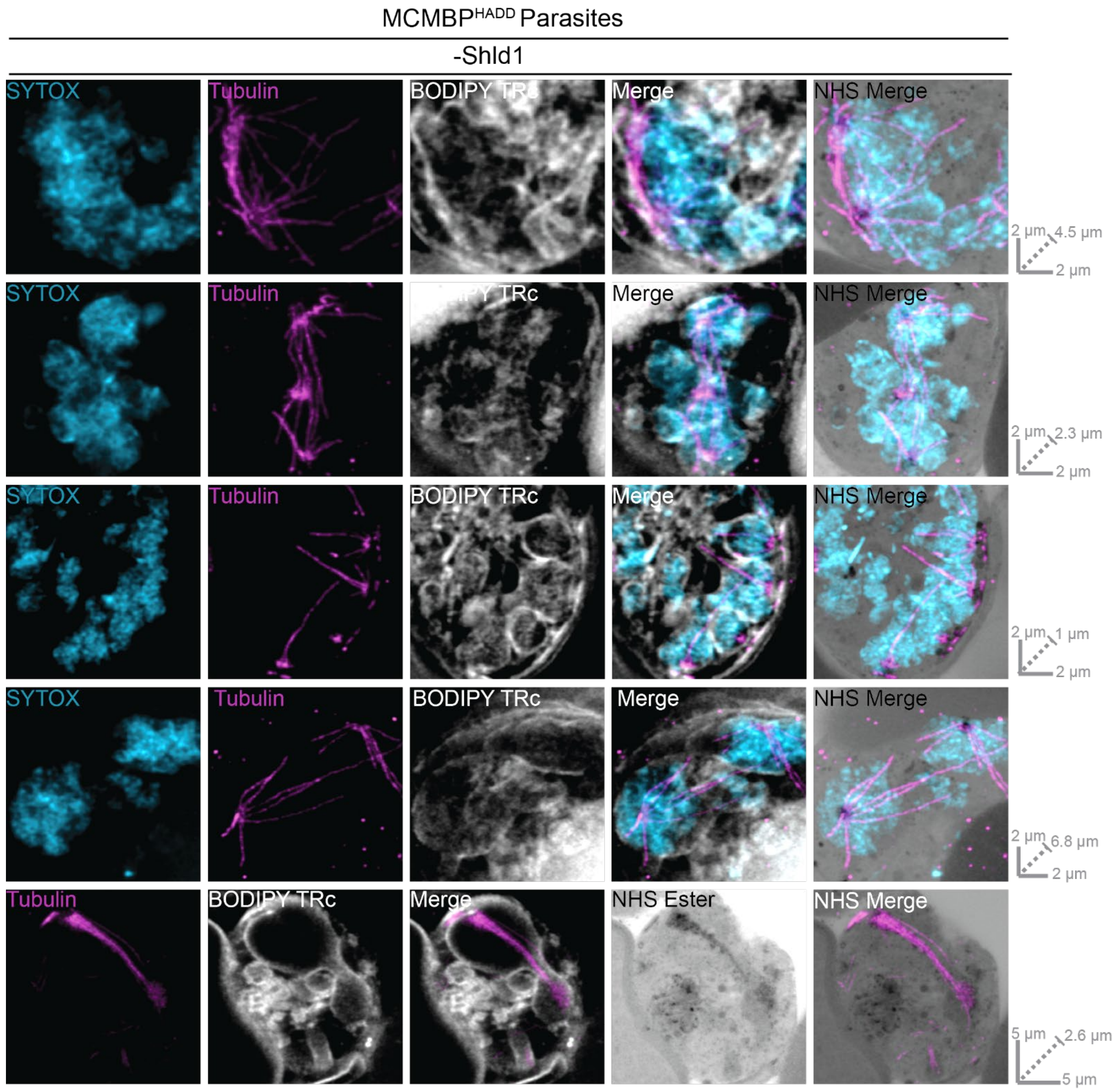


**Supplementary Figure 6. Examples of aberrant interpolar spindles, aneuploidy, and uneven DNA segregation in MCMBP deficient parasites.**
*MCMBP$^{HADD}$ parasites were cultured either in the or absence of Shld1. Parasites were then prepared for U-ExM, stained with a nuclear stain (SYTOX, in cyan), anti-tubulin (in magenta), a membrane stain (BODIPY TRc, in white), and a protein stain (NHS Ester, in grayscale), and visualized using Airyscan microscopy. Images shown here represent further examples of the defects following MCMBP knockdown described in Figure 4a. Images containing BODIPY TRc are average intensity projections, while those with NHS ester are maximum intensity projections. Slice-by-slice videos of images can be found in Supplementary Videos 15-19. Scale bars as labelled in each image, solid bars = XY scale, dashed bar = combined depth of slices used for Z-projection.*

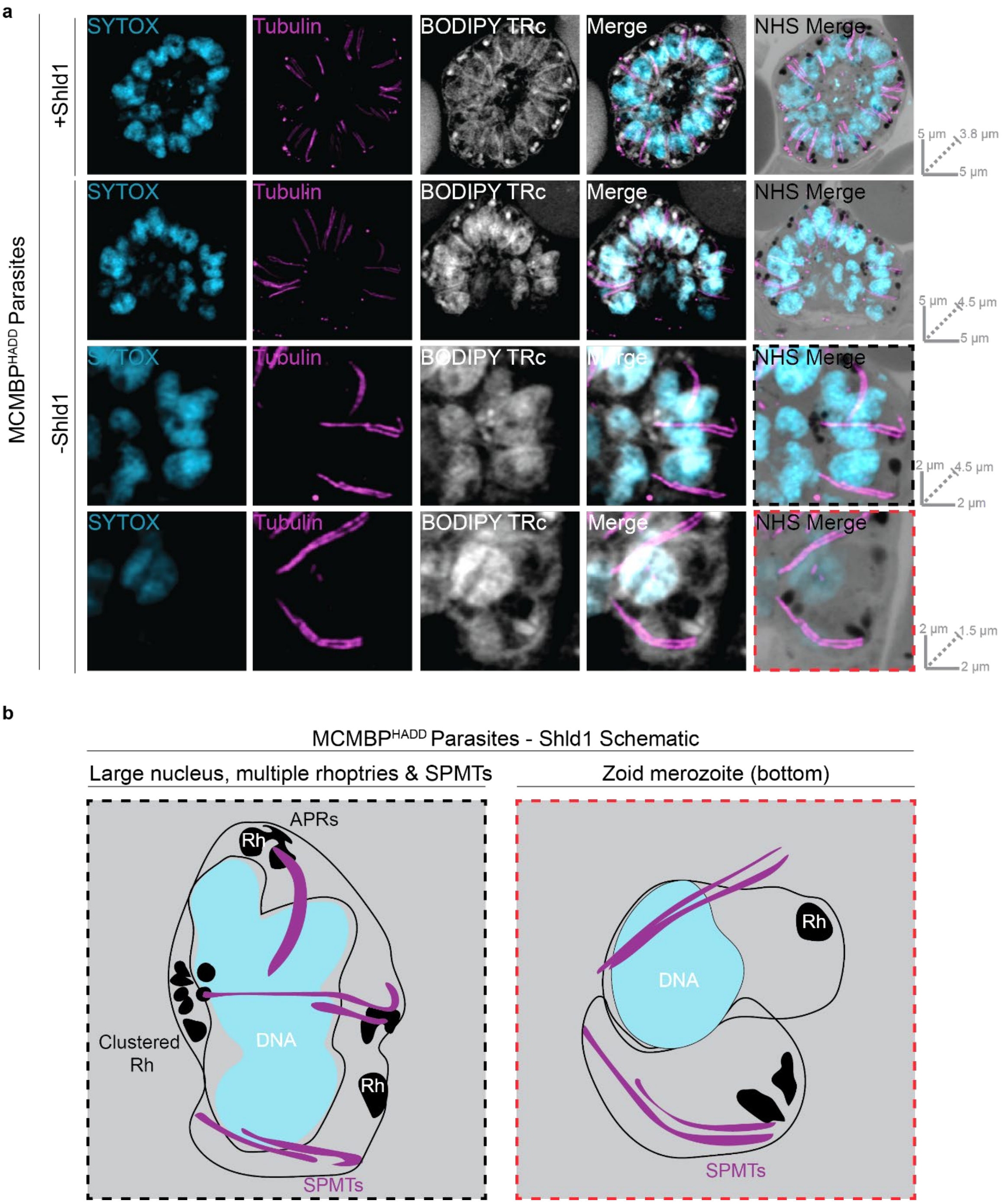


**Supplementary Figure 7. MCMBP deficient parasites display subpellicular microtubules and complete improper segmentation.**
***(a)*** *MCMBP$^{HADD}$ parasites were cultured either in the presence, or absence, of Shld1 and schizonts were arrested before egress using E64. Parasites were prepared for U-ExM, stained with a nuclear stain (SYTOX, in cyan), anti-tubulin (in magenta), a membrane stain (BODIPY TRc, in white), and a protein stain (NHS Ester, in greyscale, and visualized using Airyscan microscopy. Note that -Shld1 image rows 2 and 3 are zoomed in merozoites from the image in row 1. Images containing BODIPY TRc are average intensity projections, while those with NHS ester are maximum intensity projections. Slice-by-slice videos of images can be found in Supplementary Videos 20-21. Scale bars as labelled in each image, solid bars = XY scale, dashed bar = combined depth of slices used for Z-projection.* ***(b)*** *Schematic representations of large and zoid merozoites from* ***(a)****. Rh = Rhoptries, SPMTs = Subpellicular microtubules, APRs = Apical polar rings Black and red dashed images are the images represented in* ***(b).***
